# Effective connectivity inferred from fMRI transition dynamics during movie viewing points to a balanced reconfiguration of cortical interactions

**DOI:** 10.1101/110015

**Authors:** Matthieu Gilson, Gustavo Deco, Karl Friston, Patric Hagmann, Dante Mantini, Viviana Betti, Gian Luca Romani, Maurizio Corbetta

**Author notes:** Corresponding author; Universitat Pompeu Fabra, C/ Ramon de Trias Fargas, 25-27, 08005 Barcelona, Spain.

## Abstract

Our behavior entails a flexible and context-sensitive interplay between brain areas to integrate information according to goal-directed requirements. However, the neural mechanisms governing the entrainment of functionally specialized brain areas remain poorly understood. In particular, the question arises whether observed changes in the regional activity for different cognitive conditions are explained by modifications of the inputs to the brain or its connectivity? We observe that transitions of fMRI activity between areas convey information about the tasks performed by 19 subjects, watching a movie versus a black screen (rest). We use a model-based framework that explains this spatiotemporal functional connectivity pattern by the local variability for 66 cortical regions and the network effective connectivity between them. We find that, among the estimated model parameters, movie viewing affects to a larger extent the local activity, which we interpret as extrinsic changes related to the increased stimulus load. However, detailed changes in the effective connectivity preserve a balance in the propagating activity and select specific pathways such that high-level brain regions integrate visual and auditory information, in particular boosting the communication between the two brain hemispheres. These findings speak to a dynamic coordination underlying the functional integration in the brain.

## 1. Introduction

The brain comprises a large number of functionally distinct areas in which information and computational processes are both segregated and integrated [1, 2]. A fundamental question in system neuroscience is how information can be processed in a distributed fashion by the neuronal architecture. Brain regions exhibit a high degree of functional diversity, with a massive number of connections that coordinate their activity. Accordingly, empirical evidence from functional magnetic resonance imaging (fMRI), electro-encephalography (EEG), magneto-encephalography (MEG) in humans supports the notion that brain functions involve multiple brain areas [3]. Compared to earlier neuroimaging studies that mapped areas to function [4], the field has switched from a structure-centric viewpoint to a network-oriented one. Long-range synchronization of brain activity has been proposed as a dynamical mechanism for mediating the interactions between distant neuronal populations at the cellular level [5, 6], as well as within large-scale cortical subnetworks both at rest [7, 8, 9, 10] and when performing a task [11, 12].

Depending on the task, cortical dynamics reshape the global pattern of correlated activity observed using neuroimaging - denoted by functional connectivity (FC) [10, 13]. Presumably, both sensory-driven and cognitive-driven processes are involved in shaping FC from its resting state [14, 15]. Recently, the temporal aspect of fMRI signals has been much studied - in relation to tasks performed by subjects or their behavioral conditions - via the concept of ‘dynamic FC’ that evaluates the fluctuations of fMRI correlation patterns over time [16], the fractal aspect of fMRI time series [17, 18] or transitions of fMRI activity between areas [19]. The present study builds upon a recently developed whole-cortex dynamic model [20], which extract this functionally relevant information via the BOLD covariances evaluated with and without time shifts that describe the BOLD transition statistics.

The proposed modeling allows us to examine the respective roles played by the local variability of each brain area and long-range neuro-anatomical projections between them in shaping the cortical communication, which results in the measured FC. We rely on the well-established hypothesis that both the activity and coordination of different regions depend on both the local activity and intracortical connectivity [21]. Based on dynamic models for blood oxygen level dependent (BOLD) activity at the level of a cortical region, techniques have been developed to estimate the connectivity strengths: the notion of ‘effective connectivity’ (EC) describes causal pairwise interactions at the network level [22, 23, 24, 25]. The distinction between functional and effective connectivities is crucial here: FC is defined as the statistical dependence between distant neurophysiological activities, whereas EC is defined as the influence one neural system exerts over another [26]. In the present study, the definition of EC is actually close to its original formulation in neurophysiology [27]: estimated weights in a circuit diagram that replicate observed patterns of functional connectivity. Our model does not involve a hemodynamic response function (HRF) that maps neuronal activity to BOLD signals [28], as done with dynamic causal model (DCM) for instance [21, 29]. Nonetheless, the network EC is inferred individually for all links here (1180 connections between 66 cortical regions) and thus form a directed graph, contrasting with previous studies that directly use structural connectomes as “equivalent EC” [30, 31, 32]. For two behavioral conditions, a maximum-likelihood estimate is computed for each subject, allowing for the statistical testing of between-condition differences.

After presenting the experimental data and the whole-brain dynamic model, we reanalyze the changes observed in empirical FC between subjects at rest and watching a movie, to assess whether the proposed model is suited to capture them. Considering the estimated parameters as a signature or biomarker for the brain dynamics, we seek a mechanistic explanation for the observed FC modifications by disentangling contributions from the local variability and cortico-cortical connectivity.

## 2. Material and Methods

### 2.1. Study design for fMRI during rest and passive movie viewing

As detailed in our previous papers [33, 34], 24 right-handed young, healthy volunteers (15 females, 20-31 years old) participated in the study. They were informed about the experimental procedures, which were approved by the Ethics Committee of the Chieti University, and signed a written informed consent. Only 22 participants had recordings for both a resting state with eyes opened and a natural viewing condition; 2 subjects with only recording at rest were discarded. In the resting state, participants fixated a red target with a diameter of 0.3 visual degrees on a black screen. In the natural viewing condition, subjects watched and listened to 30 minutes of the movie ‘The Good, the Bad and the Ugly’ in a window of 24 × 10.2 visual degrees. Visual stimuli were projected on a translucent screen using an LCD projector, and viewed by the participants through a mirror tilted by 45 degrees. Auditory stimuli were delivered using MR-compatible headphones.

### 2.2. Data acquisition

Functional imaging was performed with a 3T MR scanner (Achieva; Philips Medical Systems, Best, The Netherlands) at the Institute for Advanced Biomedical Technologies in Chieti, Italy. The functional images were obtained using T2*-weighted echo-planar images (EPI) with BOLD contrast using SENSE imaging. EPIs comprised of 32 axial slices acquired in ascending order and covering the entire brain (230 x 230 in-plane matrix, IPAT = 2, TR/TE = 2 s/35 ms, flip angle = 90°, voxel size = 2.875 × 2.875 × 3.5 mm^3^). For each subject, 2 and 3 scanning runs of 10 minutes duration were acquired for resting state and natural viewing, respectively. Only the first 2 movie scans are used here, to have the same number of time points for the two conditions (i.e., 20 minutes each). Each run included 5 dummy volumes - allowing the MRI signal to reach steady state and an additional 300 functional volumes that were used for analysis. Eye position was monitored during scanning using a pupil-corneal reection system at 120 Hz (Iscan, Burlington, MA, USA). A three-dimensional high-resolution T1-weighted image, for anatomical reference, was acquired using an MP-RAGE sequence (TR/TE = 8.1 ms/3.7 ms, voxel size = 0.938 × 0.938 × 1 mm^3^) at the end of the scanning session.

### 2.3. Data processing

Data were preprocessed using SPM8 (Wellcome Department of Cognitive Neurology, London, UK) running under MATLAB (The Mathworks, Natick, MA). The preprocessing steps involved: (1) correction for slice-timing differences (2) correction of head-motion across functional images, (3) coregistration of the anatomical image and the mean functional image, and (4) spatial normalization of all images to a standard stereotaxic space (Montreal Neurological Institute, MNI) with a voxel size of 3 × 3 × 3 mm^3^. The mean frame wise displacement [35] was measured from the fMRI data to estimate head movements. They do not show any significant difference across the rest and movie recordings (*p* > 0.4): for each of the two sessions of rest, the corresponding means (std) in mm over the 22 subjects are 0.31 (0.19) and 0.31 (0.21); for movie, 0.34 (0.20) and 0.34 (0.24). Furthermore, the BOLD time series in MNI space were subjected to spatial independent component analysis (ICA) for the identification and removal of artifacts related to blood pulsation, head movement and instrumental spikes [36]. This BOLD artifact removal procedure was performed by means of the GIFT toolbox (Medical Image Analysis Lab, University of New Mexico). No global signal regression or spatial smoothing was applied. For each recording session (subject and run), we extracted the mean BOLD time series from the *N* = 66 regions of interest (ROIs) of the brain atlas used in [37]; see Table 1 for the complete list.

**Table 1:** Table of ROIs with abbreviations, names and indices. The left column indicates ensembles used later for illustration purpose, grouping ROIs into visual, auditory, so-called ‘integration’, sensory-motor, frontal and ‘central’ areas.

| Group | ROI abbreviation | Brain region | ROI index |
| --- | --- | --- | --- |
| VIS | CUN | Cuneus | 29, 38 |
|  | PCAL | Pericalcarine cortex | 28, 39 |
|  | LING | Lingual gyrus | 27, 40 |
|  | LOCC | Lateral occipital cortex | 7, 60 |
| AUD | ST | Superior temporal cortex | 14, 53 |
|  | TT | Transverse temporal cortex | 6, 61 |
|  | IT | Inferior temporal cortex | 9, 58 |
|  | MT | Middle temporal cortex | 13, 54 |
| INT | FUS | Fusiform gyrus | 5, 62 |
|  | SP | Superior parietal cortex | 8, 59 |
|  | IP | Inferior parietal cortex | 10, 57 |
|  | TP | Temporal pole | 3, 64 |
|  | SMAR | Supramarginal gyrus | 11, 56 |
|  | BSTS | Bank of the superior temporal sulcus | 12, 55 |
| SMT | PREC | Precentral gyrus | 16, 51 |
|  | PSTC | Postcentral gyrus | 15, 52 |
|  | PARC | Paracentral lobule | 30, 37 |
| FRNT | FP | Frontal pole | 4, 63 |
|  | CMF | Caudal middle frontal cortex | 17, 50 |
|  | RMF | Rostral middle frontal cortex | 20, 47 |
|  | PTRI | Pars triangularis | 19, 48 |
|  | PORB | Pars orbitalis | 21, 46 |
|  | POPE | Pars opercularis | 18, 49 |
|  | SF | Superior frontal cortex | 25, 42 |
|  | LOF | Lateral orbitofrontal cortex | 22, 45 |
|  | MOF | Medial orbitofrontal cortex | 26, 41 |
| CTRL | ENT | Entorhinal cortex | 1, 66 |
|  | PARH | Parahippocampal cortex | 2, 65 |
|  | CAC | Caudal anterior cingulate cortex | 23, 44 |
|  | RAC | Rostral anterior cingulate cortex | 24, 43 |
|  | PC | Posterior cingulate cortex | 33, 34 |
|  | ISTC | Isthmus of the cingulate cortex | 31, 36 |
|  | PCUN | Precuneus | 32, 35 |

### 2.4. Structural connectivity

Anatomical connectivity was estimated from Diffusion Spectrum Imaging (DSI) data collected in five healthy right-handed male participants [37, 24]. The gray matter was first parcellated into the *N* = 66 ROIs, using the same low-resolution atlas as with the FC analysis. For each subject, we performed white-matter tractography between pairs of ROIs to estimate a neuro-anatomical connectivity matrix. In our method, the DSI values are only used to determine the cortico-cortical skeleton: a binary matrix of structural connectivity (SC) obtained by averaging the matrices over subjects and applying a threshold for the existence of connections; see below for the optimization of the connection weights in the dynamic model. It is known that DSI underestimates inter-hemispheric connections [37]. Homotopic connections between mirrored left and right ROIs are important in order to model whole-cortex BOLD activity [32]. Here we add all such possible homotopic connections, which are tuned during the optimization as other existing connections. This slightly increases the density of structural connectivity (SC) from 27% to 28%.

### 2.5. Spatiotemporal functional connectivity from empirical fMRI data

For each session of 10 minutes (two for rest and two for movie), the BOLD time series is denoted by 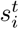 for each region 1 *≤ i ≤ N* with time indexed by 1 *≤ t ≤ T* (*T* = 300 time points at a resolution of TR = 2 seconds). We denote by 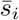 the mean signal: 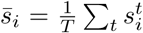 for all *i*. Following [20], the empirical FC consists of BOLD covariances without and with time shift:

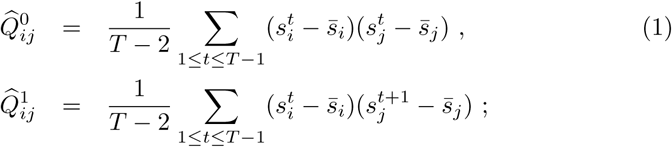

For each condition, the retained empirical FC is the mean of the two sessions. Similar calculations are done for 2 TR. BOLD correlations correspond to 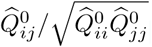. For each subject and condition, we calculate the time constant *τ*_ac_ associated with the exponential decay of the autocovariance averaged over all ROIs using time shifts from 0 to 2 TRs:

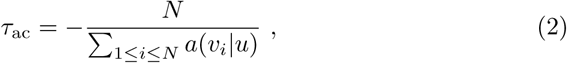

where *a*(*v*_*i*_*|u*) is the slope of the linear regression of 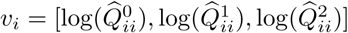 by *u* = [0, 1, 2].

### 2.6. Model of cortical dynamics

The model uses two sets of parameters to generate the spatiotemporal FC:

- the local variability embodied in the matrix Σ inputed individually to each of the *N* = 66 ROIs (see Table 1 for the complete list) or jointly to ROI pairs (only for bilateral CUN, PCAL, ST and TT);
- the network effective connectivity (EC) between these ROIs embodied by the matrix *C*, whose skeleton is determined by DSI (see details for structural connectivity above).

The rationale behind the use of spatially cross-correlated inputs (off-diagonal elements of Σ) in the model is to take into account for common sensory inputs to homotopic visual and auditory ROIs. Ideally, the model should be extended to incorporate subcortical areas and the existence of input cross-correlations inputs should be evaluated for all ROI pairs. However, this level of details is out of the scope of the present work and we constrain such input cross-correlations to 4 pairs of ROIs. Another improvement concerns the use of individualized EC skeletons for each subject or refinements of SC using graph theory for subject groups [38], but we leave this for later work.

Formally, the network model is a multivariate Ornstein-Uhlenbeck process, where the activity variable *x*_*i*_ of node *i* decays exponentially with time constant *τ*_*x*_ - estimated using Eq. (2) - and evolves depending on the activity of other populations: 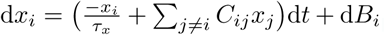. Here, d*B*_*i*_ is spatially colored noise with covariance matrix Σ, with the variances of the random fluctuations on the diagonal and cross-correlated inputs corresponding to anti-diagonal elements for CUN, PCAL, ST and TT (see Table 1). In the model, all variables *x*_*i*_ have zero mean and their spatiotemporal covariances 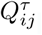, where *τ* is the time shift, can be calculated by solving the Lyapunov equation: *JQ*^0^ + *Q*^0^*J*^†^ + Σ = 0 for *τ* = 0; and *Q*^*τ*^ = *Q*^0^expm(*J* ^†^*τ*) for *τ >* 0. Here *J* is the Jacobian of the dynamical system and depends on the time constant *τ*_*x*_ and the network EC: 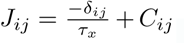, where *δ*_*ij*_ is the Kronecker delta and the superscript † denotes the matrix transpose; expm denotes the matrix exponential. In practice, we use two time shifts: *τ* = 0 on the one hand and either *τ* = 1 or 2 TR on the other hand, as this is sufficient to characterize the network parameters.

### 2.7. Parameter estimation procedure

We tune the model such that its covariance matrices *Q*^0^ and *Q*^*τ*^ reproduce the empirical FC, namely 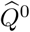 and *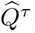*, with *τ* being either 1 or 2 TR. The uniqueness of this maximum-likelihood estimation follows from the bijective mapping from the model parameters *C* and Σ to the FC pair (*Q*^0^, *Q*^*τ*^). Apart from a refinement for the estimation of the input cross-correlation in Σ, the essential steps are similar to the iterative optimization procedure described previously [20] to tune the network parameters *C* and Σ. The model is initialized with *τ*_*x*_ = *τ*_ac_ and no connectivity *C* = 0, as well as unit variances without covariances (Σ_*ij*_ = *δ*_*ij*_). At each step, the Jacobian *J* is straightforwardly calculated from the current values of *τ*_*x*_ and *C*. Using the current Σ, the model FC matrices *Q*^0^ and *Q*^*τ*^ are then computed from the consistency equations, using the Bartels-Stewart algorithm to solve the Lyapunov equation. The difference matrices 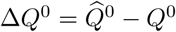 and 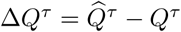 determine the model error

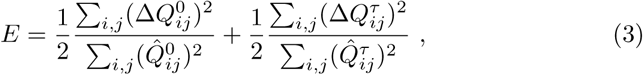

where each term - for FC0 and FC1 - is the matrix distance between the model and the data observables, normalized by the latter. The desired Jacobian update is the matrix

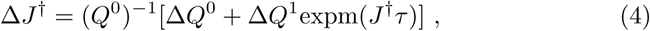

which decreases the model error *E* at each optimization step, similar to a gradient descent. The best fit corresponds to the minimum of *E*. Finally, the connectivity update is

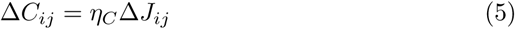

for existing connections only; other weights are forced at 0. We also impose non-negativity for the EC values during the optimization. To take properly the effect of cross-correlated inputs into account, we adjust the Σ update from the heuristic update in [20]:

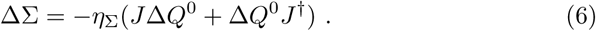

As with weights for non-existing connections, Σ elements distinct from the diagonal and cross-correlated inputs are kept equal to 0 at all times. Last, to compensate for the increase of recurrent feedback due to the updated *C*, we also tune the model time constant *τ*_*x*_ as

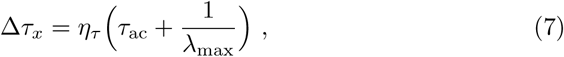

where *λ*_max_ is the maximum negative real part of the eigenvalues of the Jacobian *J*. The rationale is to avoid an explosion of the network dynamics (when *λ*_max_ → 0) while letting the model connectivity *C* develop a non-trivial structure to reproduce FC. In numerical simulations, we use *η*_*C*_ = 0.0005, *η*_Σ_ = 0.05 and *η*_*τ*_ = 0.0001.

To verify the robustness of the optimization with respect to the choice for ROIs with (spatially) cross-correlated inputs, we compare the tuned models with input cross-correlation for 1) CUN, PCAL, ST and TT; 2) CUN, PCAL, LING, LOCC, ST, TT and MT; 3) none. Although estimates differ in their detail, the results presented in this paper are qualitatively observed for all three models. In practice, the model compensates the absence of input cross-correlations by overestimating the connections between the corresponding ROIs. For simplicity, we only consider cross-hemispheric inputs for putative primary sensory ROIs involved in the task here.

A version of optimization code without cross-correlated inputs is available with the empirical data on github.com/MatthieuGilson/EC_estimation. The discarded subjects in the present study are 1, 11 and 19, among the 22 subjects available (numbered from 0 to 21).

### 2.8. Contributions of EC and input covariances in shaping spatiotemporal FC

To evaluate how the connectivity *C* and input covariances Σ contribute in shaping *Q*^0^ and *Q*^1^, we consider the models by mixing the estimates for the rest and movie conditions (with labels R and M, respectively):

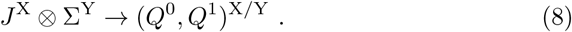

We compare them in terms of the model error defined in Eq. (3) with respect to the movie empirical FC, for which M/M is expected to give the best fit. In practice, we calculate the distance between model and empirical matrices, but considering the variances 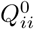 and the covariances *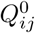* for *i ≠ j* separately within FC0; for instance, we use for variances 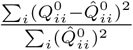, where the empirical 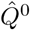 corresponds to M/M and *Q*^0^ to the evaluated model X/Y. For FC1, the matrix is taken as a whole to compute the corresponding distance.

### 2.9. Normalized statistical scores and effective drive (ED)

We define the following statistical scores for *C* or Σ to evaluate - for a given matrix element - whether the values over all subjects are consistently strong:

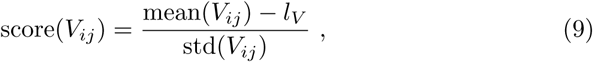

where the variable *V* is either *C* or Σ; the ‘mean’ and ‘std’ correspond to the mean and standard deviation over subjects for the considered matrix element; *l*_*V*_ is the median of all relevant non-zero matrix elements for *C* or Σ at rest, grouping the parameters over all subjects. We also define the effective drive to measure how the fluctuating activity at region *j* with amplitude corresponding to the standard deviation 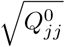 propagates to region *i*:

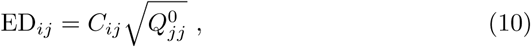

and the score(ED_*ij*_) with the corresponding median *l*_ED_ for rest.

### 2.10. Louvain community detection method

We identify communities in networks based on the modularity of a partition of the network [39]. The modularity measures the excess of connections between ROIs compared to the expected values estimated from the sum of incoming and outgoing weights for the nodes (targets and sources, respectively). The Louvain method [40] iteratively aggregates ROIs to maximize the modularity of a partition of the ROIs in community. Designed for large networks, it performs a stochastic optimization, so we repeat the detection 10 times for each subject in practice and calculate the average participation index - in the same community for each pair of ROIs over the subjects and 10 trials for each of the two conditions (rest and movie).

To test the significance of the differences between the estimated communities of each condition, we generate 1000 surrogate communities where the conditions are chosen randomly with equal chance for each subject. This gives a null distribution of 1000 participation indices for each pair of ROIs, whose upper 5% tail is used to determine significant difference across the two conditions.

## 3. Results

We start with the description of changes in the spatiotemporal FC between rest and movie, to verify if it conveys information about the behavioral condition. After ensuring that those changes are satisfactorily captured by our proposed modeling, we will examine the strongest changes in both local and network parameters (Σ and *C*, respectively) estimated across the two conditions. Compared to earlier studies with DCM that focused on changes within a specific subnetwork relying on complete Bayesian machinery [41], we infer the maximum-likelihood model parameters for each subject and seek differences from the corresponding multivariate distributions across the two conditions, as usually done with FC. Beyond individual changes for each ROI or connection, we provide an interpretation of the parameter changes in terms of activity propagation within the cortical network. Last, we will use a community analysis to detect areas with strong reciprocal activity propagation at the whole-cortex level to reach a more global viewpoint. The motivation is that - as a first guess - visual and auditory ROIs should be mainly impacted by the passive viewing task considered here [42, 43]; nonetheless, we aim to clarify whether the parameter changes imply localized or distributed effects in the cortical network [44, 12].

### 3.1. Changes in spatiotemporal FC between rest and movie viewing

We re-analyzed BOLD imaging data already reported recorded in 22 healthy volunteers when watching either a black screen - referred to as rest - or a movie (2 sessions of 10 minutes for each condition). Here these signals are aggregated according to a parcellation of *N* = 66 cortical regions or ROIs; see the list in Table 1. Firstly, we examine the changes between the two conditions in BOLD second-order statistics, which are typically used to tune whole-brain dynamic models: BOLD correlations [31, 32] and time-shifted covariances [20]. The goal is to evaluate how these observables discriminate between the two behavioral conditions. As shown in Fig. 1A, the BOLD variances (squares in the middle panel) increase by about 50% on average when watching the movie; each symbol represents a subject and the black line indicates a perfect match. In contrast, the BOLD signals do not exhibit consistent changes in their means (circles, left panel) between rest and movie. The right panel of Fig. 1A displays time constants *τ*_ac_ (triangles) estimated from BOLD autocovariance functions. They indicate the “memory depth” of the corresponding time series, quantifying how much the BOLD activity at a given time is influences by its past; see Eq. (2) in Methods. Here no significant change of temporal statistics, unlike reduction of long-range temporal correlations measured by the Hurst exponent [17]. From the plots in Fig. 1A, we discard three subjects (in red) with extreme values: two for the variances (excessive variance for movie) and one for *τ*_ac_ (small values for both conditions). From the original 22 subjects, this leaves 19 for the following analysis.

**Figure 1:**
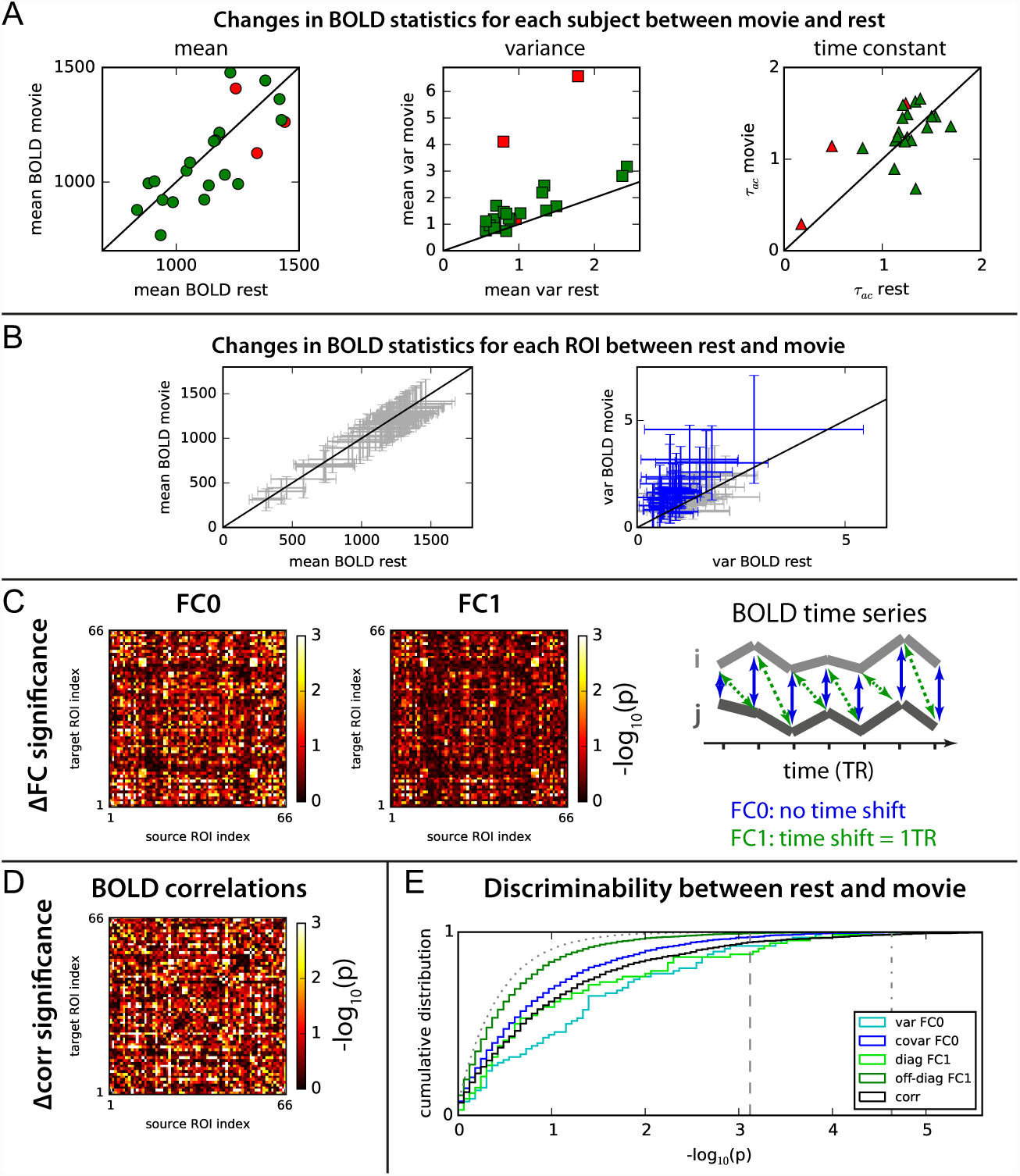
Comparison of fMRI data recorded with subjects watching a black screen (rest) or a movie. **A:** Comparison of BOLD means, variances and time constants *τ*_ac_ between the two conditions. Each symbol represents a subject and red symbols indicate the three discarded subjects, leaving 19 valid subjects for the following analysis. The black lines indicate identical values for rest and movie. **B:** Changes in BOLD means and variances between rest and movie for all ROIs. Each cross represents one of the *N* = 66 cortical ROIs and the variability corresponds to the distribution over all 19 subjects. For the variances, blue crosses indicate changes with p-value *p* < 0.01 (Welch-s t-test, uncorrected). **C:** Significance level for the changes in for all matrix elements of FC0 (no time shift) and FC1 (time shift equal to 1 TR); see Eq. (1) in Methods. For visualization purpose, an upper limit is set to 3 here, although some values are larger, cf. panel E. **D:** Same as C for BOLD correlations instead of covariances. **E:** Comparison of the cumulative distributions of p-values for variances (diagonal of FC0 in cyan), covariances (off-diagonal elements of FC0 in blue), diagonal and off-diagonal FC1 values (bright and dark green, respectively), as well as correlations (black). The dotted curve corresponds to a null distribution of p-values for two sets of 19 random variables with the same distribution. The vertical dashed line indicates the Bonferroni family-wise error rate *p* = 0.05 for *N* parameters (for the variances on the diagonal of FC0) and the dashed-dotted line for *N* (*N* − 1)/2 = 2145 parameters (for the symmetric off-diagonal covariances in FC0 and correlations).

Now considering ROIs individually and the variability of the BOLD means and variances over the subjects in Fig. 1B, we observe significant changes only for the variances in some ROIs (blue crosses). From the BOLD covariances for pairs of ROIs - both FC0 with zero time shift and FC1 with a shift of 1 TR, see Eq. (1) in Methods - we calculate the corresponding significance using Welch’s t-test (for unequal variances), which are displayed in matrix form in Fig. 1C. As a comparison, we also show the BOLD correlations in Fig. 1D: correlations seem to have overall larger values (some overlapping with FC0). The distributions of p-values for each matrix is shown in Fig. 1E, where diagonal and off-diagonal matrix elements are considered separately: all distributions exceed the null distribution (dotted dark curve) in significance. In percentage, we find more discriminative elements - with respect to the two conditions - for diagonal elements of FC1 (bright green) and variances (in cyan), followed by correlations (black), FC0 covariances (blue) and finally off-diagonal elements of FC1 (dark green). The variance-like diagonal elements of FC0 and FC1 that pass the corresponding Bonferroni threshold (dashed) correspond to 8% and 12% of all *N* = 66 elements. In comparison, less than 1% passes the corresponding threshold (dashed-dotted line for FC0 covariances and correlations), because of the large number of matrix elements; further considerations about this potential caveat are discussed later. We conclude from these results that the spatiotemporal FC defined as the BOLD covariances is informative about the behavioral condition, in line with previous studies [17, 18, 19].

### 3.2. The noise-diffusion network model captures the changes in spatiotemporal FC across conditions

In order to interpret the (significant) changes observed in the spatiotemporal FC and move beyond a phenomenological description, the present studies draws upon our recent model [20], in which the network dynamics aims to reproduce the empirical BOLD covariances, both with and without time shift. This generative model is schematically represented in Fig. 2 with only a few cortical regions in the diagrams, while the matrices involve all *N* = 66 ROIs that cover the whole cortex. Fig. 2A shows the structural connectivity (SC), which is determined by DSI data, measuring the density of white-matter fibers between the ROIs; gray pixels indicate homotopic connections that are added post-hoc, as explained in Methods. The model comprises two sets of parameters: local variability corresponding to the input covariance matrix Σ (purple fluctuating inputs in Fig. 2B) and recurrent effective connectivity (EC) between ROIs (matrix *C* with directional connections represented by the uneven red arrows). The skeleton of EC is determined by SC, assuming the existence of connections in both directions; the weights for absent connections are always zero. Here we include input cross-correlations for homotopic regions (anti-diagonal of Σ) in the visual and auditory ROIs: CUN, PCAL, ST and TT (see Table 1). The rationale is to account for binocular and binaural stimuli related to the movie stimulus; note that the corresponding parameters are estimated as others. The input structure characterized by Σ is shaped by *C* to generate the network pattern of correlated activity, which is quantified via the pair of covariance matrices, FC0 and FC1 (see Fig. 2C).

**Figure 2:**
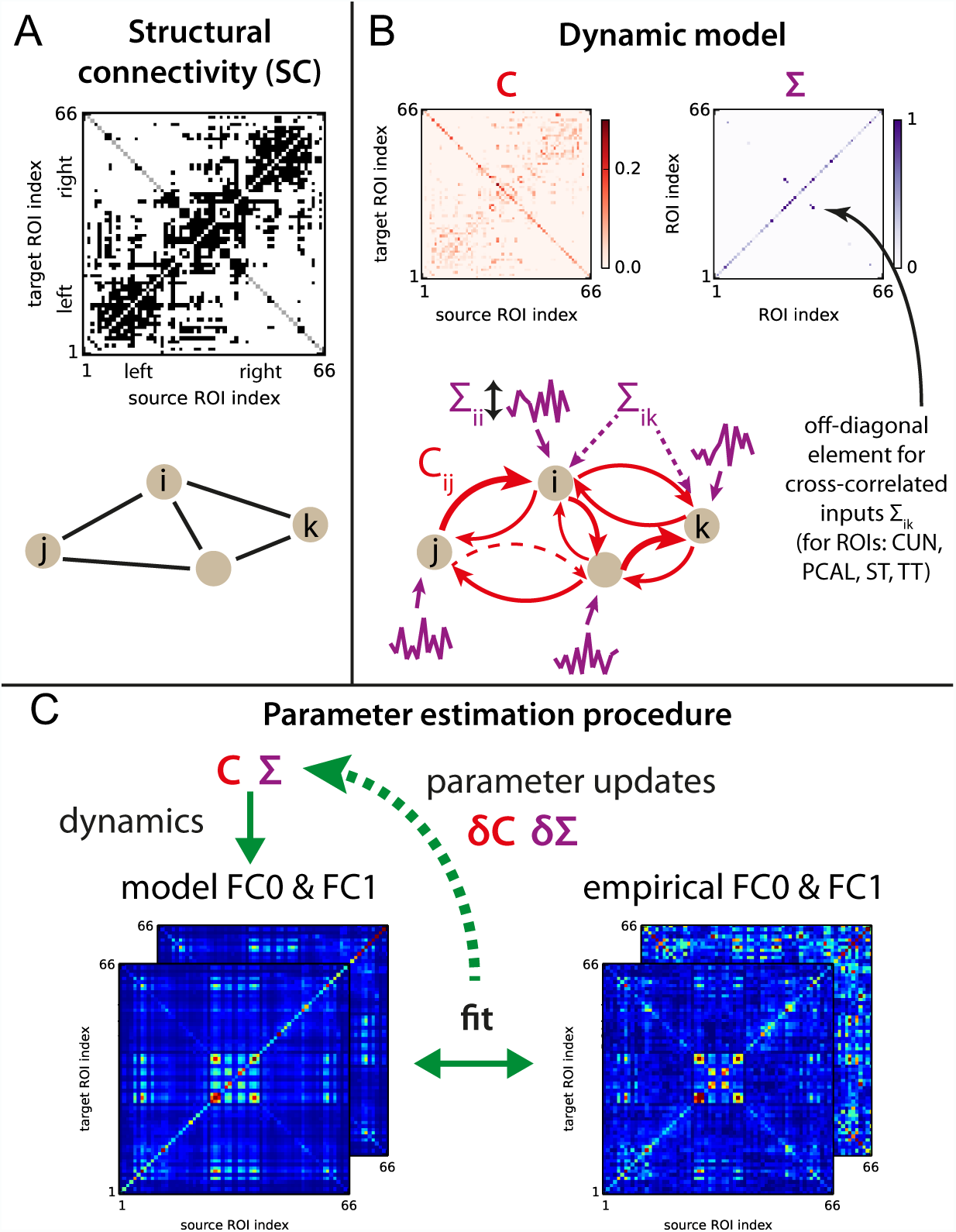
Noise-diffusion dynamic cortical model. **A:** DSI measurements provide the skeleton of the intracortical connectivity. We add inter-hemispheric connections (gray pixels on the anti-diagonal) as they are known to be missed by DSI. **B:** The parameters of the model are the recurrent effective connectivity *C* and the input covariances Σ. Contrary to SC, EC has directional connections, as represented by the red arrows with various thicknesses. Some existing connections may have zero weights (dashed arrow), equivalent to an absence of connections for the network dynamics. Here Σ comprises variances on the diagonal (one for each ROI) plus 4 pairs of symmetric elements on the anti-diagonal for cross-correlated inputs for CUN, PCAL, ST and TT (cf. Table 1). As a convention, the formatting of all matrices in this paper shows the source and target ROIs on the x-axis and y-axis, respectively. **C:** From known *C* and Σ, the model FC0 and FC1 matrices are calculated and compared to their empirical counterparts, which in turn gives the updates Δ*C* and ΔΣ for the model parameters (as well as Δ*τx*, not shown). The optimization steps are repeated until the minimal model error *E* defined in Eq. (3) is reached.

We perform the model optimization for each subject and condition. The parameters for existing connections in *C* and input (co)variances in Σ, as well as *τ*_*x*_, are iteratively tuned such that the model FC0 and FC1 best fit their empirical counterparts, as illustrated in Fig. 2C. In practice, we choose a single time constant *τ*_*x*_ for all ROIs, which is motivated by our previous results for resting-state fMRI, where no substantial difference across ROIs was observed. An improvement would consist in estimating individual values for each ROI/subject/condition, but is out of the present scope. From an initial homogeneous diagonal matrix Σ and effective connectivity *C* = 0, each optimization step aims to reduce the model error *E*, defined in Eq. (3) using the matrix distance between the model and empirical FC matrices. The best fit corresponds to the minimum of *E*, which gives the *C* and Σ estimates for each subject and condition. In summary, the model inversion explains the observed spatiotemporal FC by means of Σ and *C*.

The precision of the estimated parameters is of course limited by the number of time points in the BOLD signals, but this procedure unambiguously retrieves the model parameters for accurate empirical FC0 and FC1 observables. The iterative approach provides an advantage compared to multivariate autoregressive models applied directly to the data: it enhances the robustness of the estimation by reducing the number of estimated parameters (absent connections are kept equal to 0) and imposing constraints (non-negativity for *C* and the diagonal of Σ). Importantly, the model optimization takes network effects into account: EC weights are tuned together such that their joint update best drives the model toward the empirical FC matrices. It is also worth noting that we only retain information about the existence of connections from DSI, whose values do not influence the corresponding estimates in *C*. In practice, EC directionality strongly depends on the time-shifted covariance FC1. Further details about the model and the optimization procedure are given in Methods.

The qualitative fit of the model is displayed in Fig. 3A (left panel) for FC0 and a single subject at rest. Quantified by the Pearson correlation coefficients between the model and empirical FC matrix elements, the model goodness of fit is summarized in the right panel of Fig. 3A for all subjects and the two conditions, yielding very good values larger than 0.7 for almost all cases [32]; the model error over the subjects (mean ± std) is *E* = 0.60 ± 0.04 for rest and 0.58 ± 0.04 for movie. Importantly, we verify that the model captures the change in FC between the two conditions, as illustrated in Fig. 3B: the left panel provides the example for a subject and the right panel the summary for all subjects, as in Fig. 3A. Once again, the Pearson correlation between the model and empirical ΔFC (movie minus rest) is larger than 0.6 for most subjects. Moreover, the p-values for the changes in the FC0 matrix elements are in good agreement with their empirical counterparts in Fig. 3C, with an overall Pearson correlation of 0.37 with *p* ≪ 10^-10^. Discrepancies mainly concern absent EC connections (in black); correcting SC with the addition of missing edges may improve this aspect, but is out of the scope here. To further verify the robustness of estimated parameters, we repeat the same procedure using FC0 and FC2 with a time shift of 2 TR instead of FC0 and FC1 (with 1 TR) as done so far. We found nearly identical Σ estimates and very similar *C* estimates (Fig. 3D), which agrees with our previous results for resting-state fMRI data [20]. This confirms that the information conveyed by the transitions of BOLD activity between ROIs can be robustly extracted using the proposed dynamic model.

**Figure 3:**
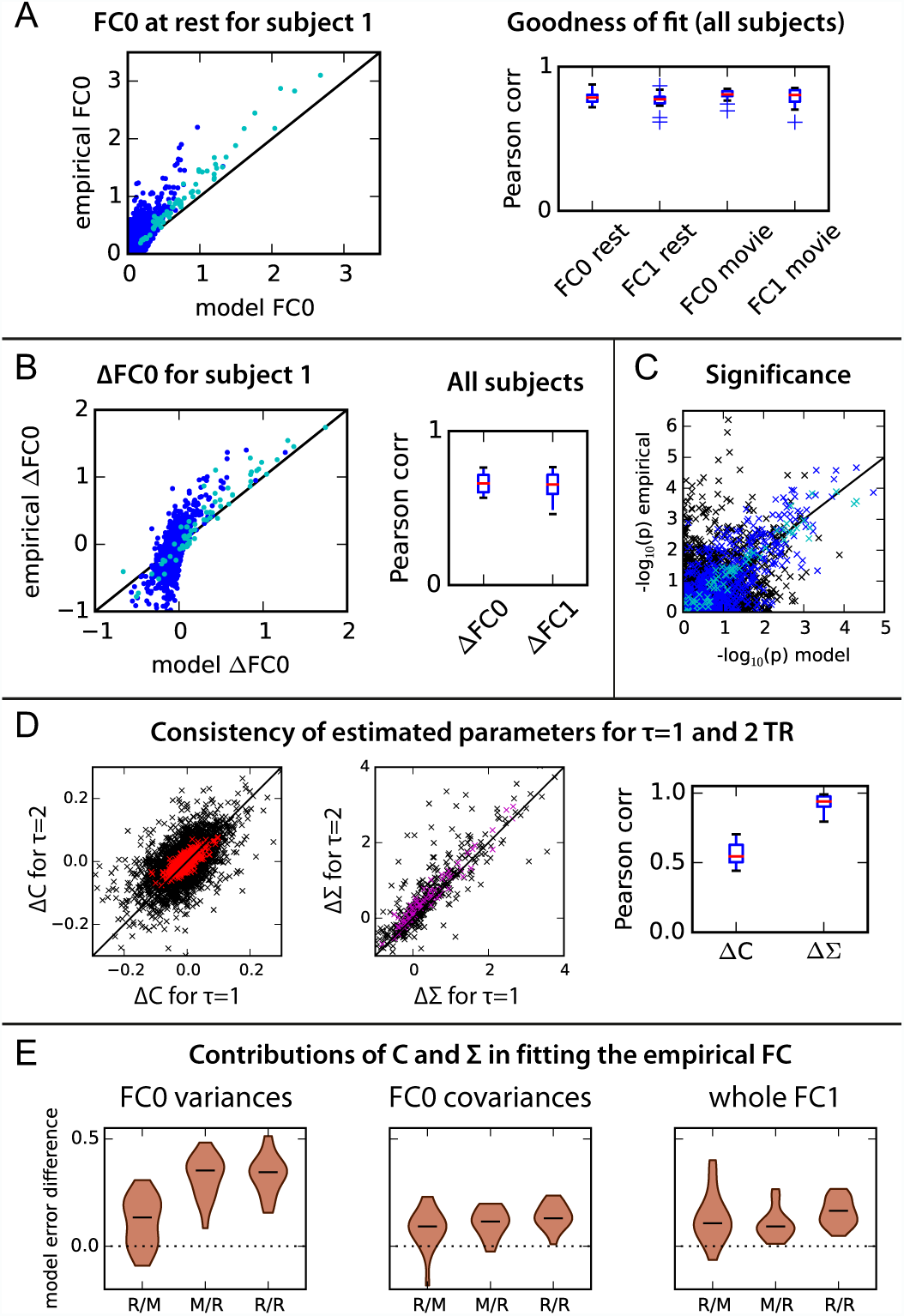
Goodness of fit of the model. **A:** The left panel shows the match between the model and empirical FC0 at rest for a single subject; each dot represents a matrix element (variances = diagonal elements in cyan, off-diagonal elements in dark blue). The right panel summarizes the goodness of fit as measured by the Pearson correlation over all subjects for the two conditions. **B:** Same as A for ΔFC0 and ΔFC1 (movie minus rest). **C:** Comparison of empirical and model p-values (Welch’s t-test) for the each matrix element of ΔFC0. Cyan crosses indicate variances, blue indicate covariances corresponding to an existing connection in EC and black covariances for absent connections. **D:** Consistency between the Δ*C* and ΔΣ matrices obtained for each subject using two distinct optimizations: FC0 and FC1 with *τ* = 1 TR; versus FC0 and FC2 with *τ* = 2 TR. The left and middle panels show the correspondence of matrix elements, with the black diagonal indicating a perfect match. Mean values over all subjects are plotted in colors. The right panel displays the Pearson correlation coefficients - one per subject - between the model estimates. **E:** Comparison of the four models combining the estimated *C* and Σ for the two conditions where X/Y corresponds to *C*^X^and Σ^Y^ with X and Y being either rest (R) or movie (M); see Eq. (8) in Methods. The violin plots indicate the difference in model error (matrix distance) between the model indicated on the x-axis and M/M, over the 19 subjects. Here the model error is decomposed according to the three components of FC: diagonal of FC0 (left panel), off-diagonal elements of FC0 (middle) and whole FC1 matrix (right).

To finally characterize how the model parameters respectively capture the FC statistics, we compare in Fig. 3E the model error for four models X/Y combining the model estimates *J* ^X^ and Σ^Y^, where X and Y are one of the two conditions, rest (R) and movie (M); see Eq. (8) in Methods for details. The M/M model is taken as a reference and others models are compared with it, in terms of error with respect the movie empirical data. The model error corresponding to the movie FC is decomposed into three components: the FC0 variances (on the matrix diagonal) and covariances (off-diagonal elements), as well as FC1 elements. As can be seen in Table 2 summarizing the p-values for these pairwise comparisons, the M/R, R/M and R/R models have significantly larger model error than M/M with *p* < 0.01 for all for all three subsets of FC; as expected, R/R consistently gives the worse fit. Changing Σ from movie to rest (M/R compared to M/M) is particularly dramatic for FC0 variances and to a lesser extent for FC0 covariances; further change *C* only affects FC1 (R/R versus M/R). Conversely, when EC is changed for rest (R/M compared to M/M), the model error increases rather homogeneously for all FC sets. However, further changing Σ for rest (R/R versus R/M) further affects FC0 elements. This illustrates that the local variability and network connectivity interplay in shaping FC and motivates a proper model inversion to interpret changes in empirical FC. In particular, a phenomenological analysis of empirical FC0 is not sufficient to estimate the change in cortical interactions (*C*) as FC0 also strongly depends on Σ, in the limit of the proposed dynamic model.

**Table 2:**
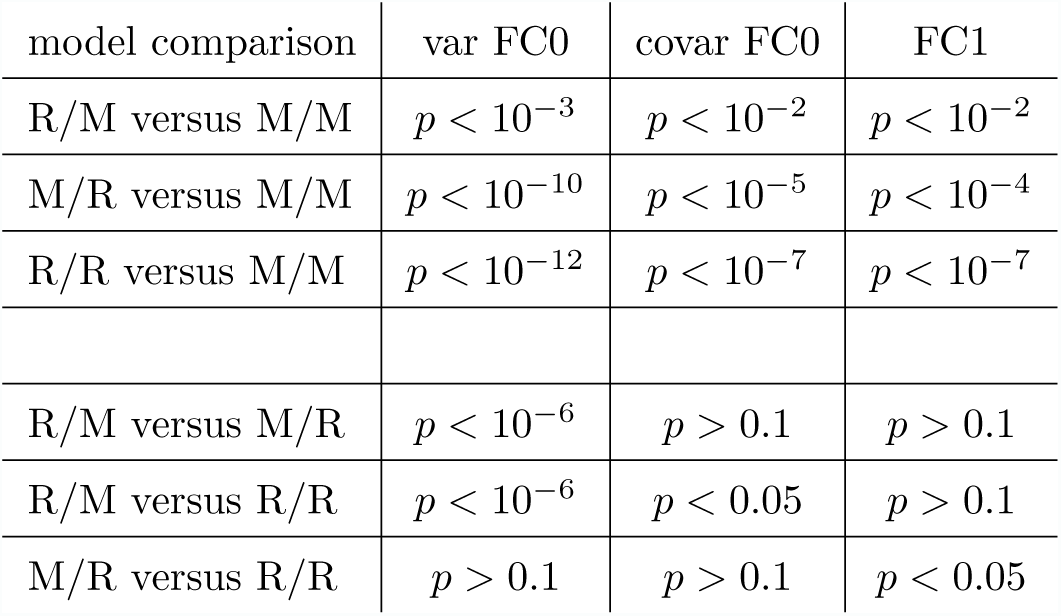
Table of p-values for pairwise comparison using Welch’s t-test between the mixed models X/Y defined in Eq. (8), as indicated on the left column. X indicates the condition for the Jacobian *J* ^X^ and Y for the input covariances Σ^Y^.

| model comparison | var FC0 | covar FC0 | FC1 |
| --- | --- | --- | --- |
| R/M versus M/M | $p < 10^{-3}$ | $p < 10^{-2}$ | $p < 10^{-2}$ |
| M/R versus M/M | $p < 10^{-10}$ | $p < 10^{-5}$ | $p < 10^{-4}$ |
| R/R versus M/M | $p < 10^{-12}$ | $p < 10^{-7}$ | $p < 10^{-7}$ |
| R/M versus M/R | $p < 10^{-6}$ | $p > 0.1$ | $p > 0.1$ |
| R/M versus R/R | $p < 10^{-6}$ | $p < 0.05$ | $p > 0.1$ |
| M/R versus R/R | $p > 0.1$ | $p > 0.1$ | $p < 0.05$ |

### 3.3. Movie viewing induces greater changes in local variability than network effective connectivity

From Fig. 3, we conclude that the tuned model satisfactorily captures the changes in empirical FC between rest and movie. Now we examine how the estimated parameters differ between the two conditions, in this way verifying whether *C* and Σ are useful signatures for changes in the cortical dynamics. We find that local variability is more affected by movie viewing than EC: at the level of the global distribution over all parameters and subjects (Fig. 4A); for the number of discriminative elements as measured by the distribution of p-values using Welch’s t-test (Fig. 4B); in magnitude as illustrated in Fig. 4C, where ΔΣ are mainly increases and Δ*C* are distributed around 0 (with a slight bias toward negative values). The Kolmogorov-Smirnov distance between the rest and movie distributions in Fig. 4A is 0.14 for Σ, to be compared with only 0.04 for *C*. Fig. 4B also shows that the outgoing EC weights (thin dark red curve, one value per ROI) experience more significant changes than individual *C* elements, but not as much as Σ elements.

**Figure 4:**
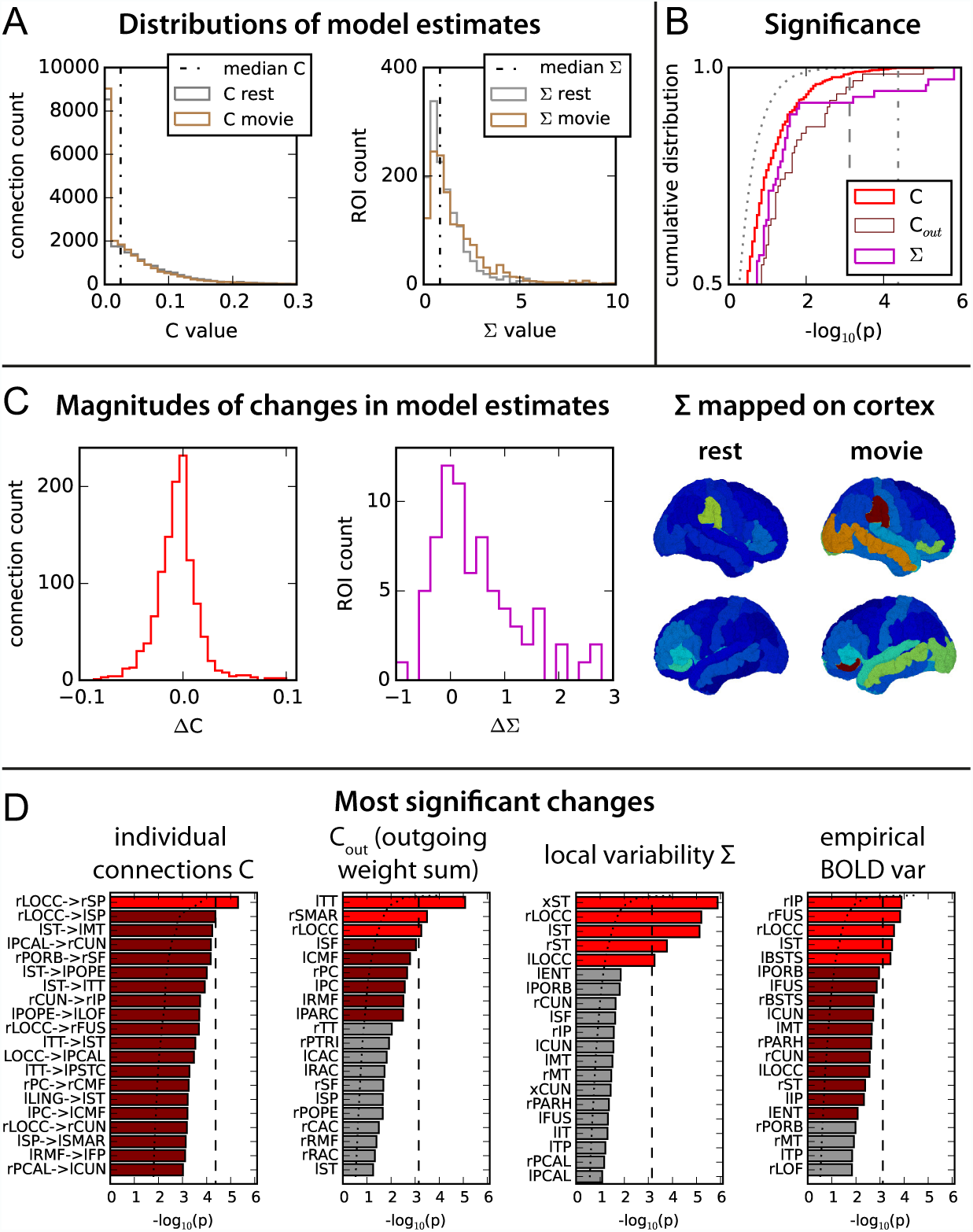
Changes in the estimated model parameters between rest and movie. **A:** Histograms of *C* and Σ values in the two conditions. The medians of the distributions for rest are indicated by the vertical dashed-dotted lines. **B:** Significance of the changes in *C* and Σ, as well as the sum of outgoing weights (*C*_out_) for each ROI. The curves correspond to the cumulative distribution over all connections/ROIs of − log_10_(*p*) for the p-value obtained from Welch’s t-test, as done in Fig. 1E for FC. The dotted curve corresponds to a null distribution of p-values for two sets of 19 random variables with the same distribution; the vertical dashed and dashed-dotted lines indicate the Bonferroni thresholds for Σ (*p* < 0.05/*m* with *m* = *N* +4 = 70 parameters) and *C* (1180 parameters). **C:** Histogram of the magnitudes of the changes - movie minus rest - Δ*C* and ΔΣ for all connections and ROI. The right panel displays the mean Σ over all subjects in each condition mapped on the cortical surface (left and right side views); hot colors indicate large values. **D:** Connections with most significant changes in *C* and ROIs with significant changes in outgoing weights *C*_out_, local variability Σ and empirical BOLD variances. For each panel, the vertical dashed line indicates the Bonferroni threshold and the dotted curve the null distribution, as in B. The ROIs passing the Bonferroni threshold are plotted in bright red, while those passing the uncorrected threshold *p* < 0.01 are in dark red. For Σ, the labels for the cross-correlated inputs to visual and auditory ROIs are indicated by ‘x’.

We now examine in Fig. 4D which connections and ROIs experience the most significant changes in the model parameters. For EC (left panel), only 1 connection in bright red passes the Bonferroni threshold with family-wise error rate equal to 0.05 (dashed line corresponding to *p* < 0.05/*m* with *m* = 1180 matrix elements), while 81 connections passed the uncorrected threshold *p* < 0.01 (7% of all connections, in dark red). These changes concern 54 ROIs among the 66; moreover, only 19 are EC increases (including the 4 passing the Bonferroni threshold) versus 55 decreases. For outgoing weights, the right TT, SMAR and LOCC show a significant increases above the Bonferroni threshold (with *m* = *N* = 66, giving 5% of all ROIs) and 6 more ROIs pass the uncorrected threshold (14%). For Σ, we identify 5 parameters that pass the Bonferroni threshold (with *m* = *N* + 4 = 70), which all concern the bilateral ST and LOCC ROIs (7% of all parameters, in bright red); no further ROI passes the uncorrected threshold. As can be seen in the right panel of Fig. 4C, the most significant changes in Σ also correspond to the largest increases, occurring for ROIs in the occipital and temporal regions.

### 3.4. Dynamical balance in the integration of visual and auditory inputs

Beyond identifying the most affected ROIs, the analysis in Fig. 4 raises the issue of comparing the significance of EC changes (Bonferroni correction with *m* = 1180 EC parameters) versus that for Σ (*m* = 70 parameters, more than one order of magnitude lower), as before with diagonal and off-diagonal of FC measures. Non-parametric permutation testing hints at the same ROIs - with slightly higher significance - but this does not solve the problem of family-wise error control. This motivates a complementary analysis focused on the network dynamics to interpret changes in *C* and Σ collectively. In particular, we examine whether the effects of Δ*C* and ΔΣ are independent of each other.

LOCC belongs to the visual cortex - even though it is not the primary visual cortex - and ST hosts the primary auditory cortex. Therefore we interpret the increase of local variability Σ for those sensory ROIs as related to the larger stimulus load in the movie viewing condition. Interestingly, changes in BOLD variances for rIP, rFUS and lBSTS that also pass the Bonferroni thershold in Fig. 4D are not straightforwardly explained by a corresponding change in the local variability Σ. This suggests that these changes could arise instead from the propagation of activity from other ROIs, as a network effect. In other words, even though changes in local activity between rest and movie are stronger than EC both in magnitude and in significance, they alone do not explain the changes for FC.

To deepen the analysis of the propagation of sensory information, we focus on the visual and auditory bilateral ROIs in our parcellation: CUN, PCAL, LOCC and LING on the one hand; ST, TT and MT on the other hand. We also consider the above-mentioned FUS and BSTS that are known to be involved in downstream visual and auditory processing [42, 43]. Fig. 5A shows the SC density between those visual (located in the lower left side and indicated by the red bars), auditory (upper right side in blue) and so-called ‘integration’ ROIs (center in purple). The dark pixels along the diagonal of the SC matrix hints at the hierarchy from visual ROIs (‘VIS’) and auditory ROIs (‘AUD’) to integration ROIs (‘INT’), corresponding to the solid arrows in the diagram on the top. In contrast, there exist fewer direct connections between VIS and AUD (dotted arrow in the diagram).

**Figure 5:**
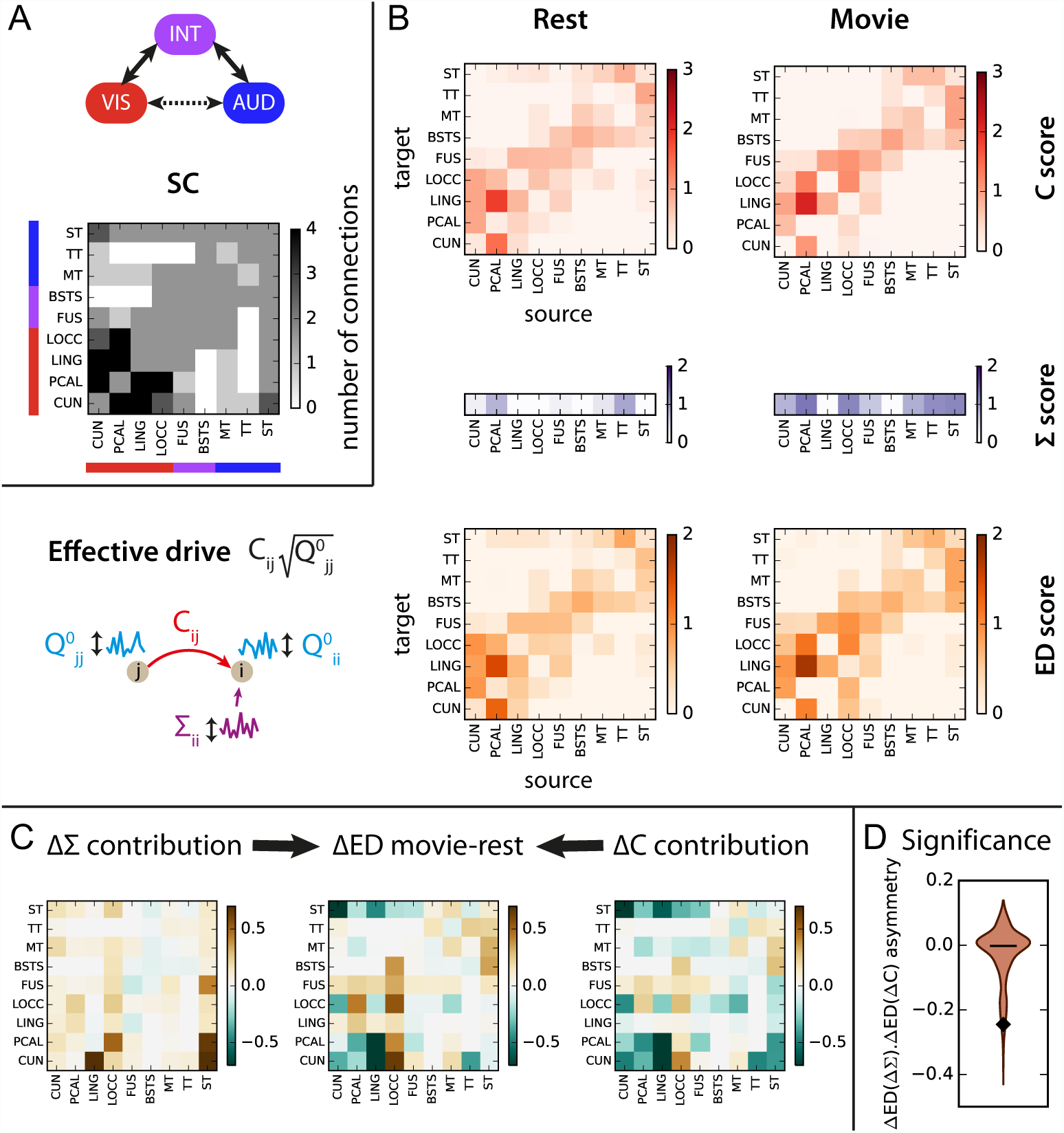
Changes in activity in the early visual and auditory pathways. **A:** Structural connectivity between 14 ROIs in the early visual (bottom left corner) and auditory (top right) pathways, as well as 4 integration ROIs (center). Connections from the left and right hemispheres are grouped together. **B:** Statistical scores for the *C*, Σ and effective drive (ED) in each condition for the ROIs and connections displayed in A. For each matrix element, the statistical score evaluates how strong the estimates are over all subjects; see Eq. (9) in Methods where the medians *l*_*V*_ for *C* and Σ correspond to the dashed-dotted lines in Fig. 4A. On the bottom left, the schematic representation describes the propagation of fluctuating activity from ROI *j* to ROI *i* quantified by ED, which contributes to the variance of the ROI activity *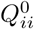* in addition to the input variance Σ_*ii*_; see Eq. (10) in Methods. **C:** Changes in ED for ROIs between rest and movie (middle panel), as well as contributions from Δ*C* (right) and ΔΣ (left). **D:** Comparison of the asymmetry of the contributions to ED from ΔΣ and Δ*C* for the subnetwork in C (diamond marker) with a null distribution obtained from similar subnetworks of randomly chosen ROIs in the rest of the network (violin plot).

The EC statistical scores in Fig. 5B - defined to identify consistently strong values over all subjects, see Eq. (9) - suggest that direct anatomical connections between VIS and AUD are not “used” for both rest and movie in practice; rather, fluctuating activity propagates between VIS and AUD via INT, back and forth. In comparison, Σ exhibits increases for most ROIs, except LING and BSTS. To further quantify the hierarchical propagation via INT, we use the effective drive (ED), which is a canonical measure for our noise-diffusion network: as illustrated in the left diagram, it measures the amount of fluctuating activity at ROI *j* sent to ROI *i* multiplied by the EC weight *C*_*ij*_ from *j* to *i*, thus contributing to *i*’s activity (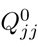 is the model variance on the diagonal of FC0, so *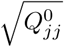* corresponds to the standard deviation). Although Fig. 5B shows a picture globally similar for rest and movie, the difference of ED scores in Fig. 5C (middle panel) indicates increases 1) from LOCC to PCAL, CUN, FUS and BSTS; 2) from almost all ROIs to FUS; 3) from ST to all ROIs except the visual ones; 4) from MT to ST and FUS. Together, most increases occur along the diagonal corresponding to the hierarchical integration mentioned above.

Thanks to our model-based approach, we can decompose the change in ED into two components related to the changes in *C* and Σ between rest and movie. If we retain only the increase in local variability Σ for the movie condition, ED increases almost everywhere, and in a particularly large amount for direct connections from ST to visual ROIs (left panel); note that visual ROIs also increase their direct effect on ST. However, negative changes in *C* nullifies this increase, as shown in the right panel. In addition, increased *C* values boost ED along the diagonal. The mixed positive and negative changes in *C* thus select pathways to preserve the hierarchical integration of sensory influx. To check whether this balancing effect is significant, we calculate the asymmetry between the left and right matrices in Fig. 5C; in practice, this is given by the scalar product of the vector obtained by stacking the matrix columns, normalized by the total ED changes in absolute value (matrix in center). The asymmetry corresponding to the 18 bilateral ROIs in Fig. 5C is represented by a diamond in Fig. 5D and compared to a surrogate distribution for the same number of randomly chosen ROIs in the rest of the cortex, while preserving the hemispheric symmetry. The significance for the observed asymmetry in the VIS-INT-AUD subnetwork is *p* < 0.04 and the 10^4^ surrogate values are mainly distributed around zero, confirming that these opposing contributions do not come artificially from the model, but from the estimated parameters instead.

### 3.5. Path selection occurs in the whole cortex

From Fig. 4, the main changes in the cortical network appear rather localized in the occipital and temporal areas. However, many “small” changes in *C* may have a more significant effect collectively and we keep in mind that the limitations of the statistical tests in the high-dimensional space of the model estimates. Therefore, we examine the changes in activity propagation at the whole-cortex level to determine whether the path selection - as an effect of the changes in the model parameters - is localized or distributed. Using the Louvain method from graph theory [39, 40], we estimate communities with higher-than-chance exchange of fluctuating activity between them, as measured by ED. We perform community analysis for each subject in each condition and pool the results over the subjects to obtain a participation index of ROI pairs, which measures the probability for them to be in the same community. To test the significance of these, we repeat the same procedure 10^3^ times while mixing the labels (rest and movie) among the subjects to generate surrogate participation indices. This gives a null distribution of 10^3^ values for each ROI pair, from which the two values for rest and movie are compared; details are provided in Methods.

Fig. 6A displays significant increases and decreases (dark and colored pixels) of participation indices for all ROI pairs in movie as compared to rest. For illustration purpose, the ROIs grouped into 6 groups: somatosensory-motor (SMT), frontal (FRNT) and so-called ‘central’ ROIs (CTRL) in addition to the visual, auditory and integration regions examined in Fig. 5; note that the integration group includes more ROIs than before, see the Table 1. Most decreases concern ROIs in the same hemisphere (bottom left and top right in the right panel), whereas increases involve interhemispheric ROI pairs (top left and bottom right in the left panel), especially ROIs from AUD, INT and FRNT. Last, interactions between VIS and INT strongly increase both within and between hemispheres.

**Figure 6:**
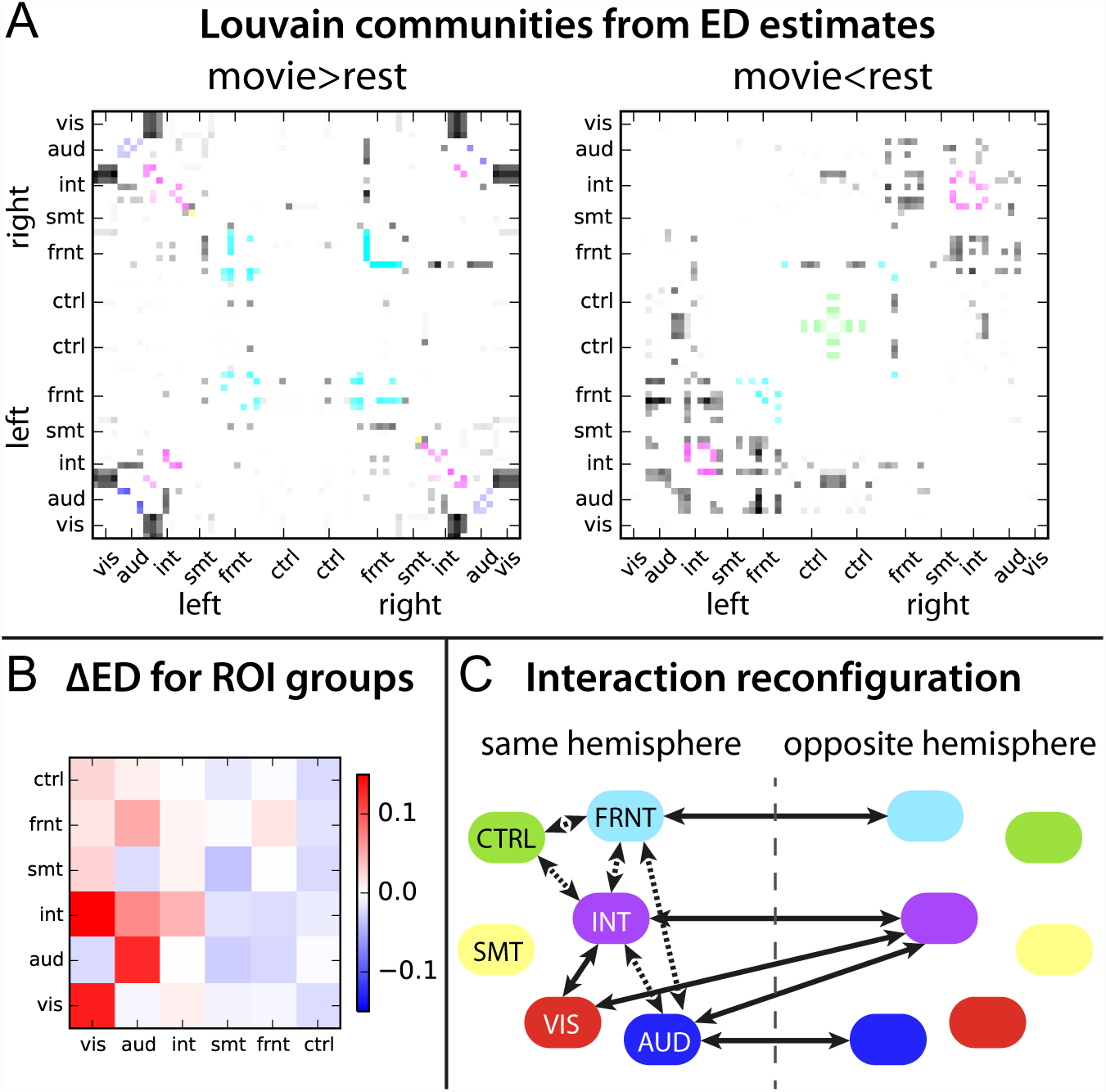
Path selection and inter-hemispheric integration in the cortex. **A:** Changes in index participation in ED-based communities estimated by the Louvain method: increases (left) and decreases (right) for movie as compared to rest. The plotted values correspond to averages over the 19 subjects, for each of which the Louvain method was applied 10 times on the ED matrix. ROIs are ordered separately for the two hemispheres according to the groups in Table 1: red for visual, blue for auditory, purple for integration, yellow for somatosensory-motor cyan for frontal and green for “central”; the pixels for ROIs belonging to the the same group are displayed in color. **B:** Changes in ED between ROIs pooled in 6 groups: visual in red, auditory in dark blue, integration in purple, somatosensory-motor in yellow, frontal in cyan and “central” (cingulate) in green. ROIs from both hemispheres are grouped together. **C:** Schematic diagram of increased interactions between ROI groups for movie compared to rest summarizing results in A and B. Arrows in solid line indicate an increase and dotted arrows decreases (sometimes mixed with increases).

To understand the changes in ED from a feedforward-feedback perspective, Fig. 6B summarizes the percentage of increase and decrease compared to rest between the groups, where the two hemispheres are taken together. The largest increases concern feedforward projections from VIS and AUD, at the noticeable exception of direct interactions between AUD and VIS. Once again, INT appears to be the intermediate that provide feedback to VIS and AUD. Fig. 6C summarizes the changes in the cortical communication, in the sense of propagation of fluctuating activity. At rest, AUD is strongly tied to the INT, SMT and part of FRNT; this cluster is decoupled in the movie condition such that part of INT can bind to VIS. Meanwhile, INT remains linked to FRNT, whose interhemispheric interactions are strongly boosted. This underlines a selective coordination of cortical paths to process sensory information in a distributed and hierarchical fashion.

## 4. Discussion

Our results shed light on a fundamental question in neuroscience: how do sensory inputs to the brain propagate via the cortical connectivity to integrate information? To address this question, we have used a recently developed model-based approach that decomposes FC in estimated parameters describing the local variability and network connectivity, EC. Moreover, the model can be interpreted in terms of cortical communication, taken as the propagation of activity across brain regions (seen via the proxy of BOLD signals). Our main finding concerns the reorganization of the cortical connectivity during movie viewing as compared to rest: although changes in EC appear at first sight smaller than those in local variability, they induce strong changes in communication across the whole cortex.

First, they are involved in a down-regulation of forward connections in a compensatory manner, such that increases in regional inputs do not saturate the network (Fig. 5D). Meanwhile, specific feedforward and backward connections are boosted to enable a hierarchical communication across ROIs, such as top-down signals from integration to sensory areas (Fig. 5C). The observed dynamic balance is expected to be task dependent - in regard with extrinsic stimulus inputs - and to result in complex patterns of functional synchronization (FC) at the network level. Our results suggest a continuum of the balanced-activity principle from the neuronal level [45] to the cortical level [46]. It is worth noting that this analysis relies on the interpretation of the brain as a dynamic system where fluctuating activity propagates; communication cannot always be simply understood at the level of single estimated parameters.

Second, the selection of specific pathways that shape the integration of sensory information acts at the level of the whole cortex. The results in Fig. 6A indicate a global reorganization of the functional communities (related to the propagation of BOLD activity) and suggest an increase of the inter-hemispheric exchange of information in the movie condition. This phenomenon is not restricted to sensory areas, but also concerns parietal and temporal areas that are related to multimodal integration [42, 43], as well as frontal areas. This highlights the need for examining the whole brain in order to understand such a distributed reconfiguration [12], extending previous studies relying on hypothesis testing for a-priori selected ROIs [47, 48, 18, 49]. The increased feedforward ED from VIS to INT illustrated in Fig. 6B is in line with explain previously reported BOLD synchronization in association areas (as well as visual areas) using inter-subject correlations (ISC) during movie viewing [44]; our estimates predict that such a synchronization should also exist in the rest of the cortex, but to a lesser extent. Moreover, our analysis hints at the same areas that were recently found to be discriminative against subjects using FC in a similar task [50]; in other words, community reconfiguration as in Fig. 6A may also be subject-specific, in addition to being task-specific.

In our study, the movie-viewing condition strongly differ from rest, as can be seen in the increased BOLD variances that presumably arise from the increase in stimulus load, in line with a previous analysis of MEG measurements for the same experiment [12]. Carefully designed experiments that controlled for the change in stimuli showed instead a decrease of variance when a subject engages a visual recognition task, as opposed to passive viewing [17]. In line with previous studies that observed a decrease in brain interactions when engaging a task [51, 49], the reconfiguration here consists in shutting down more pathways than opening new ones (cf. histogram of EC changes in Fig. 4C). This corresponds to a reduction of global synchronization (in FC), leaving aside the stimulus-related effects.

Beyond the task analyzed here, our study demonstrates that spatiotemporal BOLD (co)variances convey important information about the cognitive state of subjects, as was previously reported [17, 18]. Our spatiotemporal FC corresponds to transitions of fMRI activity at the scale of a few TRs; this statistics is averaged over the whole recording period, in contrast to other time-dependent measures such as ISC [44], metastability [52] or measures of dynamic FC averaged over 1-2 minutes, corresponding to more than 30 TRs [16]. This was already suggested by our previous analysis of resting state [20] and is in line with recent results that focused on the lag structure of BOLD signals between ROIs [53, 19]. Moreover, strong ISC during movie viewing in fMRI time series [44] - presumably arising from a locking effect to the watched naturalistic stimuli result in between-subject similarities in the corresponding covariances without time shift (FC0). The proposed framework moves beyond spatial FC (i.e., FC0 or BOLD correlations) to perform a finer analysis of fMRI measurements, in a way accounting for the effect of ISC with time shifts. Our model uses an exponential approximation of BOLD autocovariance (locally over a few TRs) and discards slow-frequency variations. In contrast, previous studies considered a broader spectrum including slow frequencies to account for long-range temporal interactions in BOLD time series [54, 17, 55]. This multifractal property of BOLD signals has been analyzed to describe undirected interactions between ROIs [18] and there is also a formal relationship between the estimation of EC described here and DCM for cross spectral density in fMRI [29]. A proper comparison with those models is left to future work.

The input “noise” related to Σ play a functional role in our model: our interpretation links the fluctuating activity (quantified by the variances in Σ) to the amount of information processed by each ROI that does not come from others. The comparison of Σ across conditions allows for a estimation of intrinsic and extrinsic components for each ROI [14, 15]. Properly modeling the fluctuating nature of brain activity [56, 57, 58] remains a challenge: models such as DCM rely on with more elaborate local dynamics to generate it [48, 29]; recent efforts aim to extend such modeling from a small number of ROIs to the whole brain [59]. Unlike DCM, we directly estimate the amplitude of random fluctuations of a simpler form of inputs (Wiener process) within each region and measure how it is transferred via the long-range projections (whose weights are estimated). In a way, our work extends analyses based on partial correlations that measure undirected interactions only [51].

Although it does not focus on the mechanisms underlying the dynamic regulation of EC [21, 48, 60], our model provides a signature of the brain dynamics that we expect to be discriminative for a broad variety of tasks and behavioral conditions. We stress again the importance of taking the whole cortex into account - or better with subcortical regions included too - to estimate such signatures. We expect a trade-off between the discriminative power and the robustness of the estimation procedure when increasing the size of the parcellation. Because *C* and Σ lie in a high dimensional space (one per connection and about one per node, respectively), statistical analysis of the estimated parameters across conditions (as well as FC-based measures) may suffer from an approach based on family-wise error correction (e.g., Bonferroni as done here). Tools from graph theory such as community analysis can be a useful complement to interpret the model estimates in a collective fashion.

## Acknowledgements

This work was supported by the Human Brain Project (grant FP7-FET-ICT-604102 to MG and GD; H2020-720270 HBP SGA1 to GD) and the Marie Sklodowska-Curie Action (grant H2020-MSCA-656547 to MG). This work was partly supported by the KU Leuven Special Research Fund (grant C16/15/070 to DM). VB was supported by a Post-Doctoral Fellowship grant from the University of Chieti.

## References

[1] D. C. Van Essen, C. H. Anderson, D. J. Felleman, Information processing in the primate visual system: an integrated systems perspective, Science 255 (1992) 419–423.

[2] B. Biswal, F. Yetkin, V. Haughton, J. Hyde, Functional connectivity in the motor cortex of resting human brain using echo-planar MRI, Magn Reson Med 34 (1995) 537–541.

[3] R. Cabeza, L. Nyberg, Imaging cognition II: An empirical review of 275 PET and fMRI studies, J Cogn Neurosci 12 (2000) 1–47.

[4] D. Cordes, V. M. Haughton, K. Arfanakis, G. J. Wendt, P. A. Turski, C. H. Moritz, M. A. Quigley, M. E. Meyerand, Mapping functionally related regions of brain with functional connectivity MR imaging, Am J Neuroradiol 21 (2000) 1636–1644.

[5] A. K. Engel, P. Fries, W. Singer, Dynamic predictions: oscillations and synchrony in top-down processing, Nat Rev Neurosci 2 (2001) 704–716.

[6] P. Fries, A mechanism for cognitive dynamics: neuronal communication through neuronal coherence, Trends Cogn Sci 9 (2005) 474–480.

[7] M. D. Greicius, B. Krasnow, A. L. Reiss, V. Menon, Functional connectivity in the resting brain: a network analysis of the default mode hypothesis, Proc Natl Acad Sci U S A 100 (2003) 253–258.

[8] M. Fox, M. Raichle, Spontaneous fluctuations in brain activity observed with functional magnetic resonance imaging, Nat Rev Neurosci 8 (2007) 700–711.

[9] M. J. Brookes, M. Woolrich, H. Luckhoo, D. Price, J. R. Hale, M. C. Stephenson, G. R. Barnes, S. M. Smith, P. G. Morris, Investigating the electrophysiological basis of resting state networks using magnetoencephalography, Proc Natl Acad Sci U S A 108 (40) (2011) 16783–16788.

[10] F. de Pasquale, S. Della Penna, O. Sporns, G. L. Romani, M. Corbetta, A dynamic core network and global efficiency in the resting human brain, Cereb Cortex 26 (2016) 4015–4033.

[11] J. F. Hipp, A. K. Engel, M. Siegel, Oscillatory synchronization in large-scale cortical networks predicts perception, Neuron 69 (2011) 387–396.

[12] V. Betti, S. Della Penna, F. de Pasquale, D. Mantini, L. Marzetti, G. L. Romani, M. Corbetta, Natural scenes viewing alters the dynamics of functional connectivity in the human brain, Neuron 79 (2013) 782–797.

[13] S. Spadone, S. Della Penna, C. Sestieri, V. Betti, A. Tosoni, M. G. Perrucci, G. L. Romani, M. Corbetta, Dynamic reorganization of human resting-state networks during visuospatial attention, Proc Natl Acad Sci U S A 112 (2015) 8112–8117.

[14] M. Mennes, C. Kelly, S. Colcombe, F. X. Castellanos, M. P. Milham, The extrinsic and intrinsic functional architectures of the human brain are not equivalent, Cereb Cortex 23 (2013) 223–229.

[15] M. W. Cole, D. S. Bassett, J. D. Power, T. S. Braver, S. E. Petersen, Intrinsic and task-evoked network architectures of the human brain, Neuron 83 (2014) 238–251.

[16] R. M. Hutchison, T. Womelsdorf, E. A. Allen, P. A. Bandettini, V. D. Calhoun, M. Corbetta, S. Della Penna, J. H. Duyn, G. H. Glover, J. Gonzalez-Castillo, D. A. Handwerker, S. Keilholz, V. Kiviniemi, D. A. Leopold, F. de Pasquale, O. Sporns, M. Walter, C. Chang, Dynamic functional connectivity: promise, issues, and interpretations, Neuroimage 80 (2013) 360–378.

[17] B. J. He, Scale-free properties of the functional magnetic resonance imaging signal during rest and task, J Neurosci 31 (2011) 13786–13795.

[18] P. Ciuciu, P. Abry, B. J. He, Interplay between functional connectivity and scale-free dynamics in intrinsic fMRI networks, Neuroimage 95 (2014) 248–263.

[19] A. Mitra, A. Z. Snyder, E. Tagliazucchi, H. Laufs, M. Raichle, Propagated infra-slow intrinsic brain activity reorganizes across wake and slow wave sleep, Elife 4.

[20] M. Gilson, R. Moreno-Bote, A. Ponce-Alvarez, P. Ritter, G. Deco, Estimation of directed effective connectivity from fMRI functional connectivity hints at asymmetries of cortical connectome, PLoS Comput Biol 12 (2016) e1004762.

[21] K. Stephan, L. Harrison, W. Penny, K. Friston, Biophysical models of fMRI responses, Curr Opin Neurol 14 (2004) 629–635.

[22] A. McIntosh, F. Gonzalez-Lima, Structural equation modeling and its application to network analysis in functional brain imaging, Human Brain Mapping 2 (1994) 2–22.

[23] K. Friston, Beyond phrenology: what can neuroimaging tell us about distributed circuitry?, Annu Rev Neurosci 25 (2002) 221–250.

[24] C. J. Honey, R. Kötter, M. Breakspear, O. Sporns, Network structure of cerebral cortex shapes functional connectivity on multiple time scales, Proc Natl Acad Sci U S A 104 (2007) 10240–10245.

[25] K. J. Friston, R. J. Dolan, Computational and dynamic models in neuroimaging, Neuroimage 52 (2010) 752–765.

[26] K. J. Friston, L. Harrison, W. Penny, Dynamic causal modelling, Neuroimage 19 (2003) 1273–1302.

[27] A. M. Aertsen, G. L. Gerstein, M. K. Habib, G. Palm, Dynamics of neuronal firing correlation: modulation of “effective connectivity”, J Neurophysiol 61 (1989) 900–917.

[28] G. M. Boynton, S. A. Engel, G. H. Glover, D. J. Heeger, Linear systems analysis of functional magnetic resonance imaging in human V1, J Neurosci 16 (1996) 4207–4221.

[29] K. J. Friston, J. Kahan, B. Biswal, A. Razi, A DCM for resting state fMRI, Neuroimage 94 (2014) 396–407. doi:10.1016/j.neuroimage.2013.12.009.

[30] J. Cabral, E. Hugues, O. Sporns, G. Deco, Role of local network oscillations in resting-state functional connectivity, Neuroimage 57 (2011) 130–139.

[31] G. Deco, A. Ponce-Alvarez, D. Mantini, G. Romani, P. Hagmann, M. Corbetta, Resting-state functional connectivity emerges from structurally and dynamically shaped slow linear fluctuations, J Neurosci 33 (2013) 11239–11252.

[32] A. Messé, D. Rudrauf, H. Benali, G. Marrelec, Relating structure and function in the human brain: relative contributions of anatomy, stationary dynamics, and non-stationarities, PLoS COmput Biol 10 (2014) e1003530.

[33] J. Hlinka, M. Palus, M. Vejmelka, D. Mantini, M. Corbetta, Functional connectivity in resting-state fMRI: is linear correlation sufficient?, Neuroimage 54 (2011) 2218–2225.

[34] D. Mantini, U. Hasson, V. Betti, M. G. Perrucci, G. L. Romani, M. Corbetta, G. A. Orban, W. Vanduffel, Interspecies activity correlations reveal functional correspondence between monkey and human brain areas, Nat Methods 9 (2012) 277–282.

[35] J. D. Power, K. A. Barnes, A. Z. Snyder, B. L. Schlaggar, S. E. Petersen, Spurious but systematic correlations in functional connectivity MRI networks arise from subject motion, Neuroimage 59 (2012) 2142–2154.

[36] J. Sui, T. Adali, G. D. Pearlson, V. D. Calhoun, An ICA-based method for the identification of optimal fMRI features and components using combined group-discriminative techniques, Neuroimage 46 (2009) 73–86.

[37] P. Hagmann, L. Cammoun, X. Gigandet, R. Meuli, C. J. Honey, V. J. Wedeen, O. Sporns, Mapping the structural core of human cerebral cortex, PLoS Biol 6 (2008) e159.

[38] V. Arsigny, P. Fillard, X. Pennec, N. Ayache, Log-euclidean metrics for fast and simple calculus on diffusion tensors, Magn Reson Med 56 (2006) 411–421.

[39] M. E. J. Newman, Modularity and community structure in networks, Proc Natl Acad Sci U S A 103 (2006) 8577–8582.

[40] V. Blondel, J.-L. Guillaume, R. Lambiotte, E. Lefebvre, Fast unfolding of communities in large networks, J Stat Mech 10 (2008) P10008.

[41] B. Li, X. Wang, S. Yao, D. Hu, K. Friston, Task-dependent modulation of effective connectivity within the default mode network, Front Psychol 3 (2012) 206. doi:10.3389/fpsyg.2012.00206.

[42] J. J. DiCarlo, D. Zoccolan, N. C. Rust, How does the brain solve visual object recognition?, Neuron 73 (2012) 415–434.

[43] T. M. Talavage, J. Gonzalez-Castillo, S. K. Scott, Auditory neuroimaging with fMRI and PET, Hear Res 307 (2014) 4–15.

[44] U. Hasson, Y. Nir, I. Levy, G. Fuhrmann, R. Malach, Intersubject synchronization of cortical activity during natural vision, Science 303 (2004) 1634–1640.

[45] N. Dehghani, A. Peyrache, B. Telenczuk, M. Le Van Quyen, E. Halgren, S. S. Cash, N. G. Hatsopoulos, A. Destexhe, Dynamic balance of excitation and inhibition in human and monkey neocortex, Sci Rep 6 (2016) 23176.

[46] G. Deco, M. Corbetta, The dynamical balance of the brain at rest, Neuroscientist 17 (2011) 107–123.

[47] R. Goebel, A. Roebroeck, D. Kim, E. Formisano, Investigating directed cortical interactions in time-resolved fMRI data using vector autoregressive modeling and Granger causality mapping, Magn Reson Imaging 21 (2003) 1251–1261.

[48] J. Daunizeau, K. Stephan, K. Friston, Stochastic dynamic causal modelling of fMRI data: should we care about neural noise?, Neuroimage 62 (2012) 464–481.

[49] R. W. Emerson, S. J. Short, W. Lin, J. H. Gilmore, W. Gao, Network-level connectivity dynamics of movie watching in 6-year-old children, Front Hum Neurosci 9 (2015) 631.

[50] T. Vanderwal, J. Eilbott, E. S. Finn, R. C. Craddock, A. Turnbull, F. X. Castellanos, Individual differences in functional connectivity during naturalistic viewing conditions, Neuroimage 157 (2017) 521–530. doi: 10.1016/j.neuroimage.2017.06.027.

[51] W. Gao, H. Zhu, K. Giovanello, W. Lin, Multivariate network-level approach to detect interactions between large-scale functional systems, Med Image Comput Comput Assist Interv 13 (2010) 298–305.

[52] G. Deco, V. Jirsa, A. McIntosh, Emerging concepts for the dynamical organization of resting-state activity in the brain, Nat Rev Neurosci 12 (2011) 43–56.

[53] A. Mitra, A. Snyder, C. Hacker, M. Raichle, Lag structure in resting-state fMRI, J Neurophysiol 111 (2014) 2374–2391.

[54] A.-M. Wink, E. Bullmore, A. Barnes, F. Bernard, J. Suckling, Monofractal and multifractal dynamics of low frequency endogenous brain oscillations in functional MRI, Hum Brain Mapp 29 (2008) 791–801.

[55] P. Ciuciu, G. Varoquaux, P. Abry, S. Sadaghiani, A. Kleinschmidt, Scale-free and multifractal time dynamics of fMRI signals during rest and task, Front Physiol 3 (2012) 186.

[56] R. B. Stein, E. R. Gossen, K. E. Jones, Neuronal variability: noise or part of the signal?, Nat Rev Neurosci 6 (2005) 389–397.

[57] A. A. Faisal, L. P. J. Selen, D. M. Wolpert, Noise in the nervous system, Nat Rev Neurosci 9 (2008) 292–303.

[58] I. Dinstein, D. J. Heeger, M. Behrmann, Neural variability: friend or foe?, Trends Cogn Sci 19 (2015) 322–328.

[59] S. Frässle, E. I. Lomakina, A. Razi, K. J. Friston, J. M. Buhmann, K. E. Stephan, Regression DCM for fMRI, Neuroimage 155 (2017) 406–421. doi: 10.1016/j.neuroimage.2017.02.090.

[60] D. Battaglia, A. Witt, F. Wolf, T. Geisel, Dynamic effective connectivity of inter-areal brain circuits, PLoS Comput Biol 8 (2012) e1002438.

